# WeightedLD: The Application of Sequence Weights to Linkage Disequilibrium

**DOI:** 10.1101/2021.06.04.447093

**Authors:** Oscar J. Charles, Joseph Roberts, Judith Breuer, Richard A. Goldstein

## Abstract

Sequence-weighting methods are commonly employed to account for biases in sequence datasets. We use a weighting scheme which considers the observed distinctiveness of sequences and apply it to calculations of linkage disequilibrium. Each sequence now contributes a weighted score to linkage disequilibrium measurements of pairwise loci. We demonstrate that this reduces the effect of uneven sampling, as underrepresented groups of sequences will each contribute more individually than redundant, similar sequences.

**Availability:** Source code for a python and rust implementation are freely available at under an MIT license at github.com/ojcharles/WeightedLD.

## Introduction

Assembling a set of sequence data is a common first step in population genetics projects. The composition of such sets is affected by factors such as geography, accessibility of different populations, similarity to laboratory strains and the idiosyncratic nature of sequencing efforts. This results in sequencing data that is highly biased, complicating and compromising downstream analyses.

Sequence weighting schemes have been developed to compensate for overrepresentation of populations in genetic data. The more redundant and less distinct a sequence is, the lower its weight and the less it contributes to downstream analyses. Conversely a sequence that is divergent will be assigned a relatively high weight and will more strongly influence later calculations. While there is consensus that sequence weighting is important, there are a number of different methodologies ^1,2^.

Linkage Disequilibrium (LD) measures the non-random association between genetic markers across different sites in the genome ^3^. LD analyses are frequently applied to pairwise variable loci in a multiple sequence alignment (MSA) in order to test for the presence of recombination. Alleles presenting long-range LD can suggest functional similarities of encoding gene products ^4^, the presence of population admixtures, epistatic selection, or other selection pressures ^5^.

Despite its widespread use in sequence analysis, sequence weighting has not yet been applied to most population genetics calculations such as LD. Consider a set of sequences representing disproportionately sampled populations, by geography or clinical presentation. It is possible that noteworthy signals represented in the minor populations, and/or interplay between the major and minor populations would be diluted by the dominant linkage of alleles in the major population. We describe how sequence weighting can alleviate much of the effect of biases in sample composition in calculations of LD. This represents a general methodology which could be applied to a wide range of population genetic studies.

## Methodology

Consider a multiple sequence alignment derived from one or more specific populations. The composition of such an alignment has inevitable biases, such as an overrepresentation of samples from golf-playing nations (GPNs) or nearly exact copies of laboratory strains. We wish to calculate LD in a manner that is robust to these compositional biases.

We define *p_i_* to be the frequency of allele *A_i_* at a locus *q_j_* to be the frequency of allele *B_j_* at a second locus, and *g_ij_* as the joint frequency of sequences with both alleles at these loci. if there is complete random association between these two loci, *g_ij_* = *p_i_ q_i_*. This will not be true in the presence of linkage disequilibrium. The amount of the discrepancy is quantified by *D*, the linkage disequilibrium coefficient, given by ^6,7^.

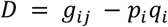

As *D* will be affected by the amount of variation at these two sites, it is common for this quantity to be normalised

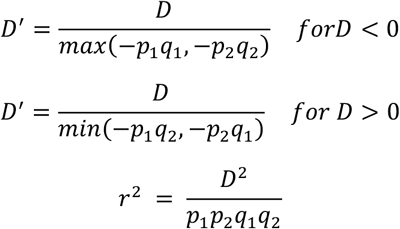

Next, we introduce sequence weights into this framework. If the weight of sequence *l* is *w_l_* and we are considering two sites, one where sequence *l* has allele *x*_1_, the other where it has allele y_1_, then:

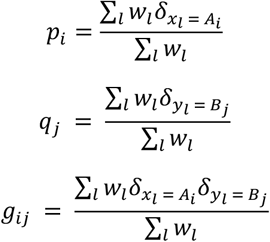

For calculating these weights, we use the approach developed by Henikoff and Henikoff ^8^.

## Implementation and testing

We implemented a procedure that calculates LD using sequence weighting and the LD scores above in both Python and Rust ^9,10^. The software accepts a multiple sequence alignment or variant call file as input, identifies all variable positions and then returns a matrix of values for [position *i*, position *j, D, D′, r*^2^] per pairwise comparison. To account for ambiguous bases there is an option to set --min-acgt fraction, which will throw out variable positions which contain below this fraction of A, C, G, or T. Also, there is a --min-minor argument, which ignores positions with a (dominant) minor fraction less than the specified value. We benchmarked the rust implementation compute time with default settings on an i7-8650U laptop with 32GB RAM. Sequence data were obtained from NCBI or the 1000 genomes project and aligned with MAFFT ^11–13^. The Hepatitis C weighting and LD calculations shown below completed in 0.35 seconds, 100000 Sars-cov2 global whole genomes from 2021 in 1.02s and Human GRch38 chromosome 19 positions 44000000-45000000 from 200 diploid individuals, evenly split among GBR and PJL populations required 21.53s.

Sequence weighted LD calculations should be beneficial in reconstructing a more accurate population LD when only biased data is available. To demonstrate this we generated an unbiased MSA containing two sets of 137 whole genome sequences of Hepatitis C subtype 1 of Japanese and USA origin. This alignment was unevenly sampled to create a second biased alignment of 25 Japanese and 137 USA sequences. With these two alignments we computed three sets of results using WeightedLD, 1) Unbiased-unweighted 2) Biased-unweighted and 3) Biased-Weighted. Figures 1 and 2 demonstrate that assigning weights to sequences results in higher contribution from the minor Japanese population, shifting LD pairwise calculations towards the values obtained with an unbiased set. The following results were calculated using parameters --min-acgt 0.9 --min-minor 0.02 --r2-threshold 0.02. Within the figures the Wilcoxon test, Pearson correlation coefficients, and graphics were generated in R ^14^.

**Figure 1.**
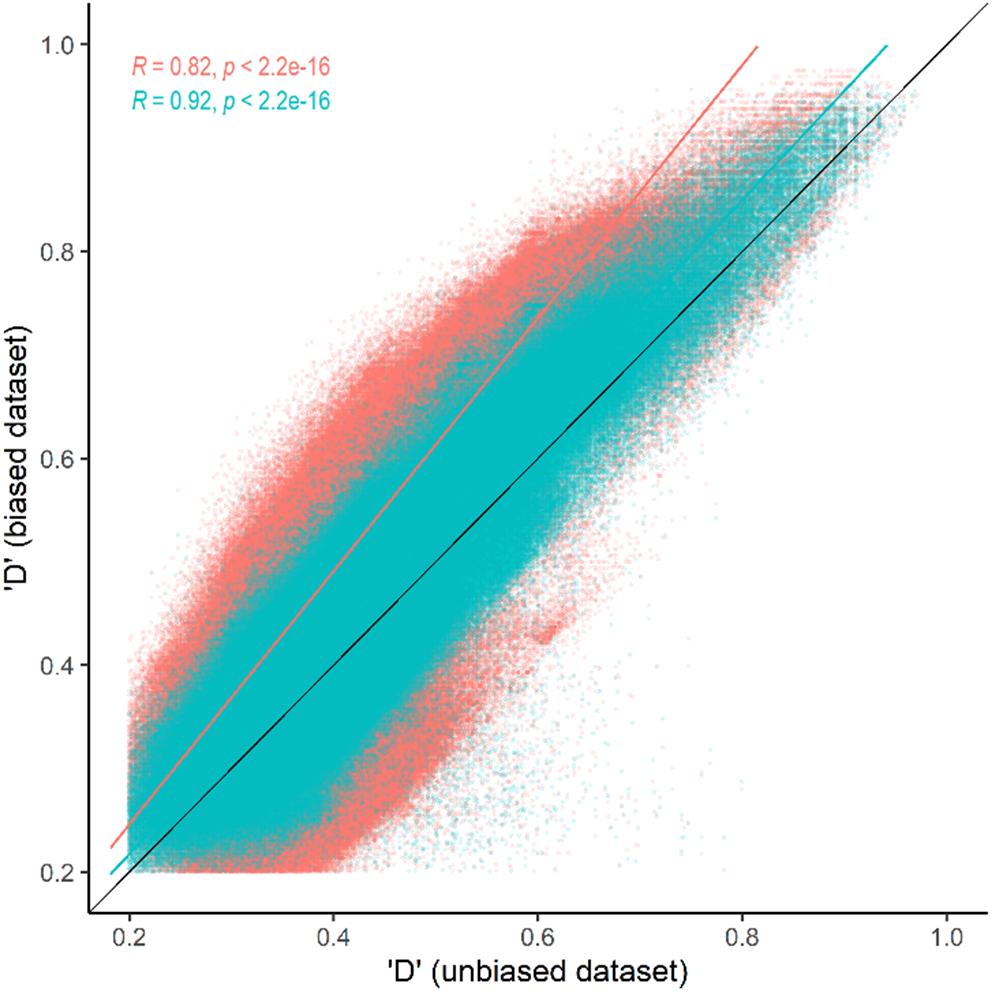
Weighted D’ values of each pairwise comparison of variable sites obtained for the biased unweighted (vermillion) and biased weighted (blue) datasets compared with that obtained with the unbiased dataset. For each point the error is represented by the distance from the black central line.

**Figure 2.**
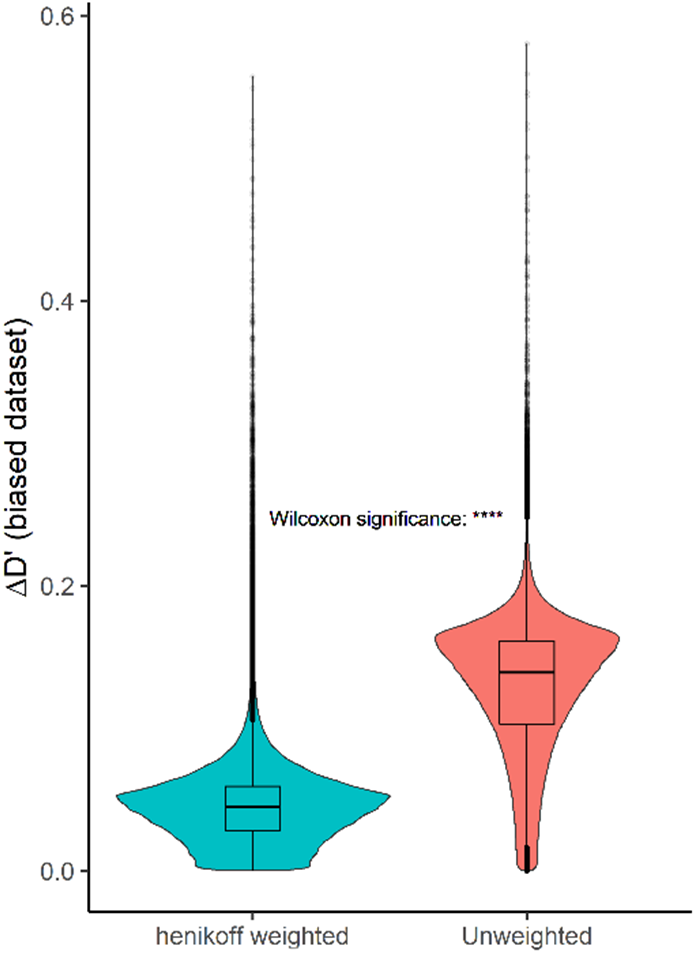
Violin plot of the absolute differences in LD D’ of the biased unweighted dataset (vermillion) and biased weighted dataset (blue) relative to that obtained with the unbiased sample. Mean ΔD’ for unweighted and Henikoff weighted were 0.129 and 0.045 respectively.

